# Semantic Tuning of Single Neurons in the Human Medial Temporal Lobe

**DOI:** 10.1101/2025.10.21.682935

**Authors:** Katharina Karkowski, Marcel S. Kehl, Yingjie Qin, Alana Darcher, Philipp Karkowski, Valeri Borger, Rainer Surges, Martin N. Hebart, Thomas P. Reber, Stanislas Dehaene, Yair Lakretz, Florian Mormann

## Abstract

The *Medial Temporal Lobe* (MTL) is key to human cognition, supporting memory, emotional processing, navigation, and semantic coding. Rare direct human MTL recordings revealed concept cells, which were proposed as episodic memory building blocks due to their context- and modality-invariant response. However, there is a long-standing debate whether concept cells encode information in a discrete, all-or-none fashion, or as a graded, continuous function of semantic dimensions. To resolve this, we developed a closed-loop protocol that analyzes neuronal spiking in real time and adaptively presents new stimuli based on semantic similarity to response-eliciting previous ones. We found that human concept cells show graded responses, following *semantic tuning curves*. Furthermore, the tuning width and steepness of MTL neurons vary, with the hippocampus exhibiting the steepest tuning. Regional analysis of these tuning curves yields region-specific categorical response profiles (e.g., food responses in the amygdala). Finally, we show that at the population level, MTL activity is best captured by continuous semantic vector models, which outperform categorical and network-based metrics. Together, these findings establish that concept cells exhibit semantic tuning curves, providing new mechanistic insights into the neural organization of semantic information in the human brain.

## Introduction

The human mind has the capacity to construct concepts – abstract representations of the surrounding world ^1–4^ – by mapping objects and recurring patterns in the environment onto stable, discrete entities. This process of structuring knowledge supports many aspects of higher cognitive functions, such as flexible categorization, thought, reasoning, and long-term memory.

The human MTL – a core region for declarative memory formation^5,6^ – contains a distinct type of neurons that respond to a given concept regardless of context and invariant to the sensory modality through which the concept is presented ^7–9^. These neurons, also known as concept cells, have been hypothesized to serve as building blocks for declarative memory by encoding the meaning of a concept ^10^, a hypothesis only recently confirmed experimentally^11^. Recordings of individual neurons in the human MTL have provided fundamental insights into the function of these concept cells. They respond to various images of the same person as well as to their written and spoken name, indicating an amodal representation of their identity^7^. Neurons have also been found that are tuned to abstract entities like numbers ^12^ or to the semantic categories of objects ^13^ such as clothes ^14^, animals ^15^, or scenes ^16^. Remarkably, some MTL concept neurons have been shown to even extend their semantic tuning to odors ^9^. Yet the fundamental coding principles that govern their selectivity, invariance, and generalization have remained a matter of debate. Some research suggests that these neurons respond in a binary fashion ^17^, while other studies indicate that the strength of their response varies continuously with the semantic content of the stimuli^14,18^. A recent study further proposes that these neurons respond to regions within a high-level visual feature space ^19^.

Here, we investigated single-cell activity in the human MTL (Figs. 1a&b) and found that the activity of concept cells can be characterized by *semantic tuning curves*. We show that concept cells respond to stimuli as a continuous sigmoidal function of their semantic similarity to a peak in conceptual space — much like the tuning curves identified in visual and auditory modalities to orientation and frequency, respectively, but tuned to a neuron’s preferred semantic content. We characterize the semantic fields that drive concept-cell activation, the width and steepness of their tuning (Fig. 1c), and the semantic categories and features that trigger these cells across individual MTL regions. Furthermore, we use these data to test which theory of semantic coding – network-based, feature-based, or continuous-vector semantics – best aligns with the human MTL encoding. Finally, we investigate the existence of cells with multiple activation peaks. Together, this research demonstrates that semantic tuning curves are a fundamental feature of conceptual representation, providing a new mechanistic perspective to understand how the brain supports complex cognitive abilities such as memory formation and abstract thought.

**Fig. 1:**
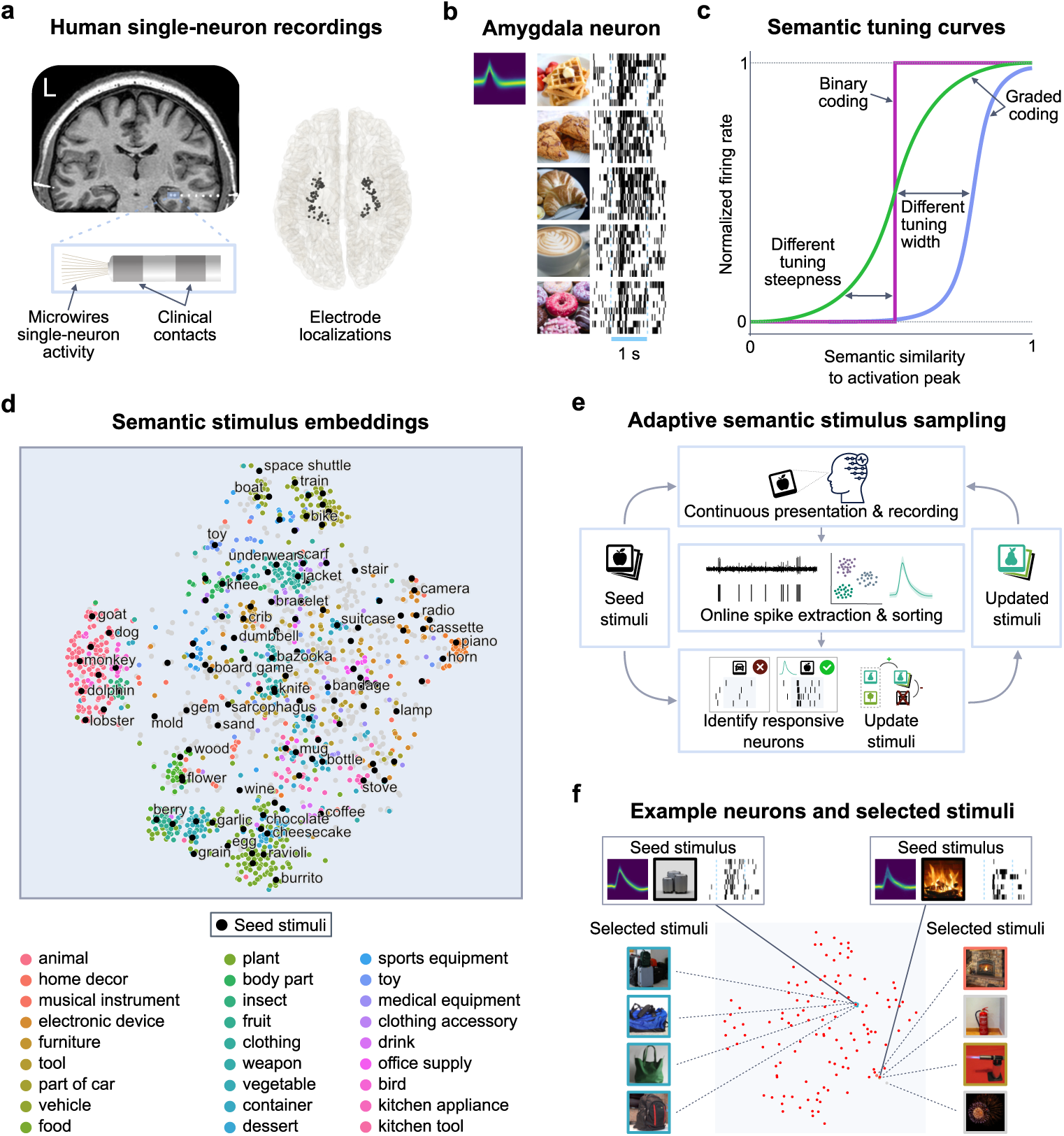
General experiment setup. a) Single neuron recording setup. The figure shows the micro-electrode locations of all patients for all electrodes included in the study. b) Spiking activity of neuron #60 (amygdala) responding to different food items. Each row shows a stimulus alongside the corresponding spiking activity rasters. c) Hypothetical tuning curves. The curves represent competing hypotheses about the tuning of concept cells ranging from a binary tuning to a smooth transition and also varying tuning widths. d) Concepts of the THINGS database. The concepts are embedded in a 2D-space using *t-distributed stochastic neighbor embedding* (t-SNE), colored by category. Initial stimuli for the paradigm are marked in black. e) Experiment structure: A set of 120 ’seed stimuli’ is presented to map the initial semantic space. Based on a neuron’s real-time spiking activity, ’updated stimuli’ are then automatically selected and presented. f) Stimuli updating mechanism: New stimuli are selected in the neighborhood around response-eliciting stimuli. Neuron #249 (entorhinal cortex) responded to *suitcases* and neuron #235 (amygdala) to *fire*.

## Results

All experiments were conducted in patients with pharmacologically intractable epilepsy undergoing chronic seizure monitoring at the Department of Epileptology at the University Hospital Bonn. Patients were implanted with clinical depth electrodes to identify the seizure focus for potential surgical treatment, as well as high-impedance microwires. From these microwires, we recorded a total of 2, 835 neurons across the anatomical target regions (amygdala, hippocampus, entorhinal cortex and parahippocampus; Hereafter, A, H, EC, PHC, respectively). Of these, 333 were characterized as single units that responded to at least one concept (Fig. A.1a).

### Closed-Loop Real-Time Selection of Semantically Related Stimuli

In order to study the semantic receptive fields of concept cells, we developed an experimental paradigm that evaluates neural data in real time. Once a putative concept cell is identified, together with its associated response-eliciting stimulus, our approach allows to automatically select new concepts with a controlled degree of semantic similarity to the original stimulus and to present them within the same experimental recording session. This approach allows for a fine-grained investigation of the semantic tuning curves of concept cells, which would be difficult to achieve with predefined stimuli given the vast, potentially infinite space of pictures and concepts and the limited patient time in a clinical setting.

We focused on concrete concepts that can be unambiguously represented by images from the THINGS database ^20^, which contains a wide range of everyday objects. Each concept is represented by a set of images depicting it in natural contexts, accompanied by a semantic embedding that serves as a metric of semantic relatedness between concepts ^21^. Fig. 1d shows a 2D-visualization of the concepts in semantic space. Semantically similar items are located close to each other, and concepts in the same category are mostly clustered together. For example, animals can be found on the middle left, while vehicles are located at the top, musical instruments on the right, and food-related items at the bottom.

The core of the paradigm is a closed-loop algorithm that controls which images are shown during subsequent trials based on neural responses from preceding trials (Fig. 1e). Specifically, the paradigm starts by displaying an initial set of seed images and performs real-time spike sorting to find responses of MTL neurons, which are used to automatically select subsequent stimuli (Fig. 1e). To ensure we only include responses to semantic content rather than visual features, a different image representing the same concept was chosen for each trial. An example of the paradigm may look as follows: If images of a suitcase elicit strong activity in a neuron, the paradigm will pick semantically similar objects, like duffel bag and backpack, for subsequent trials (Fig. 1f). As stated, this approach allows for a fine-grained investigation of concept cell semantic tuning curves that is challenging with predefined stimuli, given the vast space of concepts and limited patient time.

### Semantic Tuning Curves in Concept Neurons

To understand how MTL neurons respond to stimuli and how semantic similarity affects their spiking activity, we created heatmaps of the firing rates of neurons throughout the experiment. Figs. 2a&b show two examples of such heatmaps for neurons from the amygdala and hippocampus, respectively. Cold colors indicate low firing rates, while hot colors represent high firing rates across the full semantic space of the THINGS dataset (Fig. 1d). For visualization purposes, the semantic space is projected onto two-dimensions with interpolated response magnitudes (RM).

**Fig. 2:**
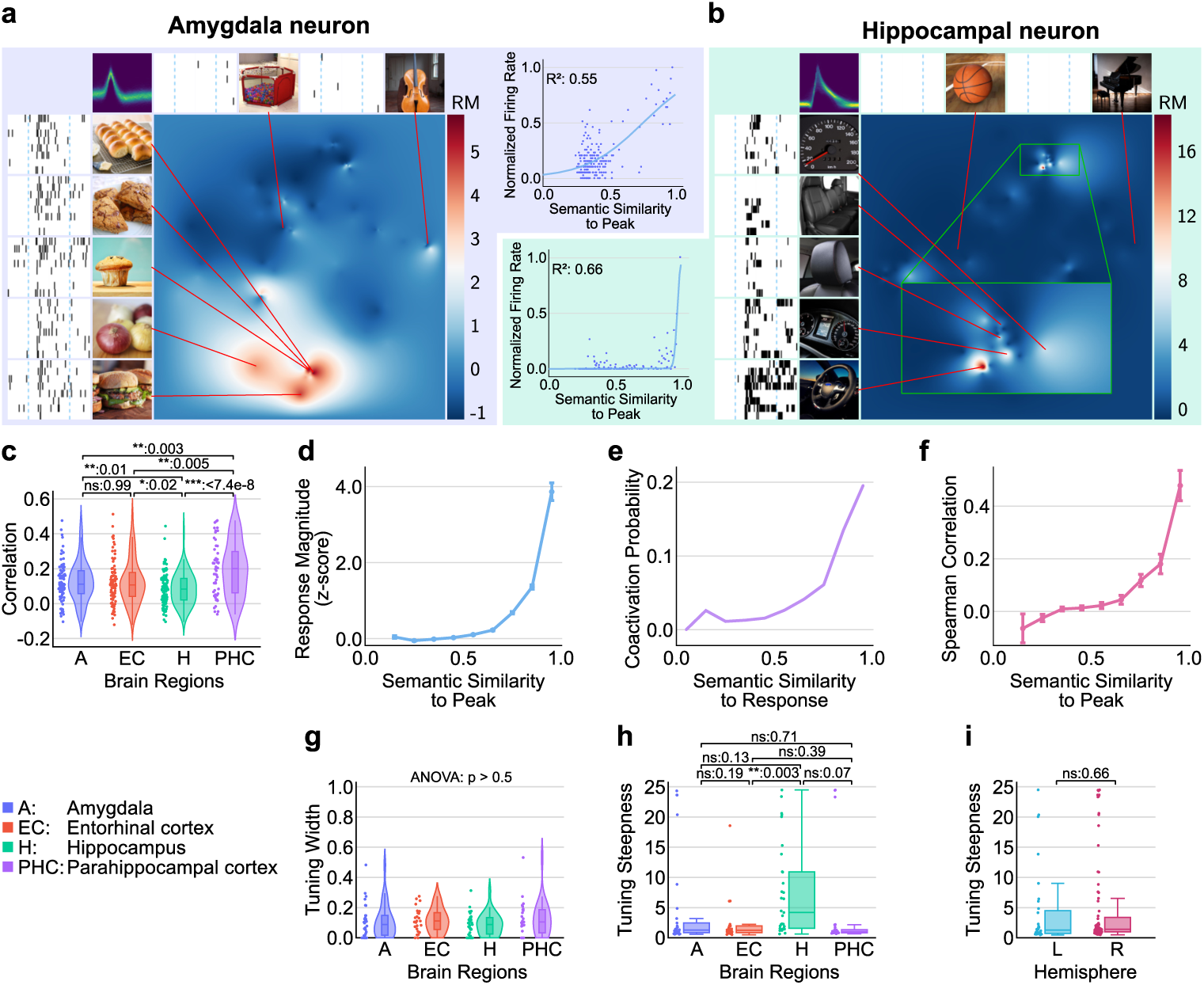
Characteristics of semantic tuning curves in the human MTL. a) Left: Semantic tuning heatmap interpolated over the response magnitude (RM) for an example neuron (Neuron #75, amygdala). This neuron shows strong activity for bakery items and minimal response to other objects like a violin. Right (top): Graded response strength based on semantic similarity for the same neuron and logistic curve fit with *R*^2^ = 0.55. b) Equivalent to a, but for a second example neuron (Neuron #191, hippocampus). c) Spearman correlation between similarities in semantic space and neural activation for different MTL regions. d) Averaged response magnitudes across all recorded neural data reveal a semantic tuning curve with a transition between low and high activation. e) Probability that an average neuron responds to two stimuli as a function of their semantic similarity. Given a response-eliciting stimulus, the likelihood of another stimulus evoking a response in the same neuron decreases with increasing semantic distance in space. f) Spearman correlations between firing rate and semantic similarity, calculated within discrete semantic similarity bins. Correlations are high for stimuli within the activation peak’s vicinity and become uncorrelated outside of the neuron’s semantic receptive field. g) Distributions of semantic tuning widths for different MTL regions show considerable variability among neurons, with a median width of around 0.1 (IQR=0.13). h) Regional distributions of tuning curve steepness across all neurons, indicating a relatively smooth increase in semantic tuning, with a median steepness of 1.27 (IQR=2.89). i) Comparison of tuning curve steepness between hemispheres. c,g,h,i) All boxes depict the median and interquartile range.

The example amygdala neuron (Fig. 2a) exhibited a pronounced increase in activity in response to several food items. Responses were confined to concepts in a specific region of the semantic space, with little or no neural activity elsewhere. Critically, the neuron’s response within this food-related region was also graded, depending on the specific food item presented; for example, bakery items caused more activity than onions. In contrast, less semantically related stimuli, such as a violin, elicited much weaker or no spiking activity in this neuron. This pattern of selective and graded responsiveness across semantic categories and within an active region is what we refer to as a neuron’s *semantic tuning curve* (see further examples in Fig. A.2). Similarly, Fig. 2b presents a hippocampal neuron with a distinct semantic tuning curve; this neuron showed strong activity for stimuli related to car parts (e.g., car seats, dashboard) but not for others, such as a piano. Notably, the active semantic region for this neuron was more narrowly confined than that of the neuron in Fig. 2a.

To quantify the semantic tuning curves of concept neurons and their properties, we modeled their response profiles using logistic regression. For this, we first identified the activity peak for each neuron based on the ’center of mass’ of the most response-eliciting stimuli (*Methods*). This yielded a vector for the peak, which describes the point in semantic space of maximal activity – we refer to this vector as the *peak embedding*. Next, for each neuron, we computed the semantic similarity between all other stimuli and the peak embedding and, finally, fit a logistic function between the neuron’s firing rate and semantic similarity to the peak (*Methods*). We chose a logistic function because its two parameters directly represent the width and steepness of the semantic tuning curve (Fig. 1c). This allowed us to quantify the variability of these properties across all neurons in our dataset, as illustrated by two example neurons. Figs. 2a (top right) and 2b (bottom left) also show the resulting fitted tuning curves for our two example neurons, demonstrating that the second neuron (‘car parts’) indeed has a narrower width and a steeper slope compared to the first neuron (‘food-related items’).

### Semantic Tuning Across the Neural Population

Having demonstrated semantic tuning in individual example neurons, we next sought to systematically quantify the relationship between firing rates and semantic similarity of concepts across the entire population of recorded concept cells. To this end, we first computed the Spearman correlation between each neuron’s normalized firing rate and the semantic similarity to its activation peak. Fig. 2c illustrates the distributions of these correlations across all neurons, consistently showing correlations significantly greater than zero across anatomical target region (*t >* 8.5, *p <* 1×10^−10^ for one-sample t-tests against zero for each region), with the strongest mean correlation observed in the parahippocampal cortex. No significant differences were found between the left and right brain hemisphere (*p* = 0.72, Fig. A.1b). Averaging the response magnitudes across all recorded neurons revealed a semantic tuning curve with a continuous transition between low and high activation (Fig. 2d). This result was also robust for alternative metrics for semantic relatedness (Fig. A.1). Complementing this, we found that the probability of a neuron responding to two stimuli depended on their semantic similarity. Given a response-eliciting stimulus, the likelihood of another stimulus evoking a response in the same neuron decreased with increasing semantic distance (Fig. 2e). Finally, a close inspection of single-neuron tuning curves (Figs. 2a top right and 2b bottom left) also explains why the overall Spearman correlation in Fig. 2c might seem relatively low: We observed that when computing the correlation in different similarity ranges (Fig. 2f), this correlation primarily held for stimuli within the vicinity of the activation peak, while firing rates were uncorrelated outside the neuron’s specific semantic receptive field.

Having established the graded nature of semantic tuning, we next sought to characterize its specific parameters. For this, we examined the distributions of fitted parameters from the logistic models (Fig. 1c, Figs. 2a top right and 2b bottom left), focusing on neurons where the model provided a relatively good fit (*R*^2^ *>* 0.1).

We defined a neuron’s tuning width as 1 minus its inflection point (the point of maximum steepness of the logistic fit, which corresponds to the threshold level of semantic similarity above which the neuron’s firing rate exceeds 50% of its peak-to-trough range). Fig. 2g shows the distributions of tuning-curve widths split by regions of the MTL, demonstrating that tuning width varied considerably among neurons. The median tuning width was 0.1 (*IQR* = 0.13). For example, concept pairs with a semantic distance of 0.1 include ”peach/orange,” ”penguin/dolphin,” and ”doughnut/cookie.” The widest tuning observed was around 0.4, exemplified by pairs like ”rattlesnake/kiwi,” ”caviar/candy,” and ”shoe/crown.” This suggests that the semantic field of MTL neurons typically covers a relatively limited region of semantic space rather than a broad expanse. We found no significant differences in tuning width across different brain regions (Fig. 2g) or between hemispheres (Fig. A.1c). The steepness of the tuning curve is reflected by the slope at the inflection point. Across all neurons, the median steepness was 1.27 (*IQR* = 2.89), indicating a relatively smooth increase in the semantic tuning curve. For instance, the tuning curve shown in Fig. 2a has a tuning width of 0.25 and a steepness of 1.15.

Interestingly, the hippocampus exhibited markedly steeper tuning curves compared to other regions (Fig. 2h), suggesting that hippocampal neurons often have tuning curves that more closely resemble a step function (*p <* 0.005 for a one-sided t-test comparing hippocampus to all other regions). However, it is important to note that even a very steep tuning does not automatically imply binary coding. Fig. 2b shows an example of a neuron in the hippocampus, demonstrating steep yet graded tuning. Similar to tuning width, there was no significant difference in tuning steepness between hemispheres (Fig. 2i).

Earlier work has suggested a binary coding in the human MTL^17^. However, when reanalyzing our data for such binary coding, we observed clear graded responses, most pronounced once the normalization stimulus was removed (Fig. A.5).

In sum, these results suggest that MTL concept neurons exhibit *semantic tuning curves*, with firing patterns confined to specific parts of the entire semantic space. Within their preferred semantic domain, activity was found to be graded, with the neuron’s firing rate varying depending on how semantically similar a given item is to the peak response.

### MTL Regions Encode Distinct Semantic Domains

Beyond individual neurons, we next quantified the semantic preferences of the neural population of each MTL region (A, EC, H and PHC). To visualize each region’s preferred semantic fields, we first averaged the z-scored heatmaps (e.g., Fig. 2a) across all neurons (Fig. 3a). The amygdala, for instance, showed a preference for food items, while the parahippocampal cortex favored vehicles and animals. In contrast, the entorhinal cortex and hippocampus exhibited a more uniform coverage of semantic space, lacking strong preferences for specific semantic domains.

**Fig. 3:**
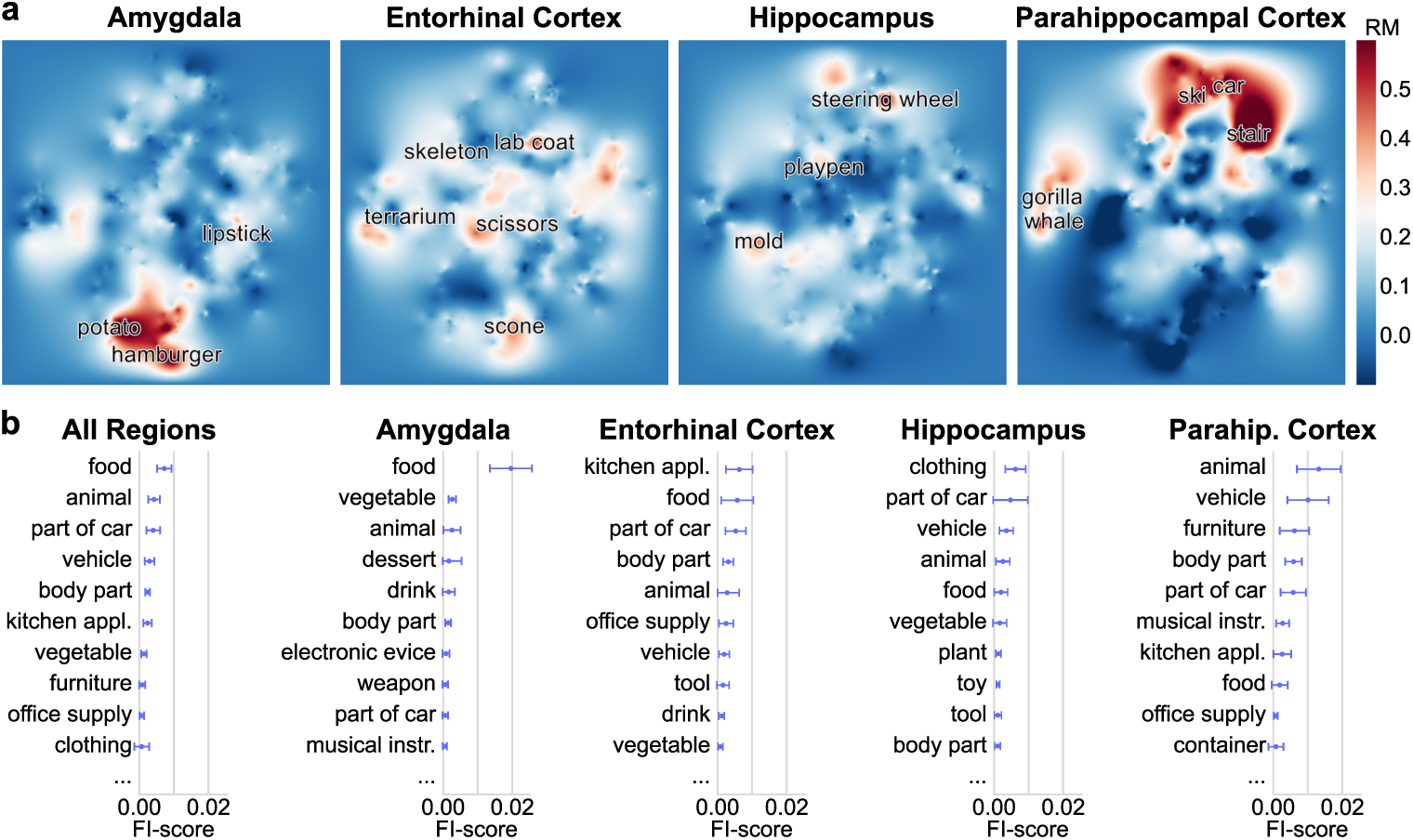
Semantic categories driving the activity of MTL neurons. a) Heatmaps of interpolated response magnitudes (RM), averaged across all neurons within each brain region. b) Top 10 categories by mean FI score, averaged across neurons for different brain regions.

To capture the semantic preferences observed in the heatmaps in more interpretable terms, we mapped the firing patterns to well-defined semantic categories available in the THINGS dataset. While heatmaps offer a detailed view of responses across the entire semantic space, directly associating them with categories concisely describes a neuron’s or a region’s overall preferred semantic field. This does not imply strict, all-or-none selectivity, but rather aims to capture the general semantic domain that elicits the strongest or most consistent activity.

The THINGS dataset provides a list of high-level, superordinate categories for each object. For example, the object ‘cookie’ belongs to the ‘food’ and ‘dessert’ category. To identify which semantic categories best explain a concept cell’s spiking activity, we fit a regression model for each cell to predict this activity from semantic categories (see Fig. A.3 for the correlations among all categories in our dataset). To estimate each category’s individual contribution to predicting spiking activity, we then computed feature importances (FIs; *Methods*) for all categories. FIs measure how strongly a semantic category predicts spiking activity. For example, the FI scores for the two example neurons in Fig. 2a,b quantitatively confirm selective tuning to food- and car-related objects, respectively (see Fig. A.4). Fig. 3b shows the mean FI-score for all categories across all MTL neurons. Categories with the highest average scores include food, animal, part of car, and vehicle, whereas some other categories have average FI-scores close to zero. These analyses reveal differences between MTL brain regions, consistent with the heatmap visualizations in Fig. 3a. For example, neurons recorded in the amygdala showed a clear preference for food and food-related categories. Although the highest average scores in the entorhinal cortex were for ’kitchen appliance’ and ’food’, and for ’clothing’ and ’part of car’ in the hippocampus, both regions demonstrated weaker specificity to semantic categories overall. The parahippocampal cortex showed a preference for animals and vehicles, possibly due to a general preference for outdoor objects ^16^.

Together, these results demonstrate that different MTL regions exhibit distinct preference profiles for semantic domains, with the amygdala and parahippocampal cortex showing the most prominent preference for specific stimulus categories.

### Which Semantic Theory Best Accounts for Neural Encoding of Concepts in the Medial Temporal Lobe?

How concepts are organized in the human mind is a central and long-standing question in cognitive science. Here, we tested how well various theoretical models of concept organization in the mind aligned with the firing patterns of MTL neurons. We contrasted three theoretical approaches to semantic similarity (Fig. 4a): (1) *network-based models*, (2) *feature-based models*, and (3) *continuous-vector models* ^22^. Network-based models represent words as individual nodes in a large graph, where edges connect closely related words, expressing for instance hierarchy (as between *apple - fruit*, *tree - plant* )^23,24^. Semantic similarity is given by the length of the path between two concepts in the network. Feature-based theories assume that words are represented in memory as a set of discrete features (e.g., ’has a tail’, ’has wings’) ^25,26^. Semantic similarity can then be determined based on the number of shared features between two concepts ^25,27^. Lastly, continuous-vector models represent concepts as vectors in a high-dimensional space, which may be derived from corpora, for example, using co-occurrence statistics, following the distributional hypothesis ^28–30^, or based on behavioral experiments, such as an odd-one-out task^31^.

**Fig. 4:**
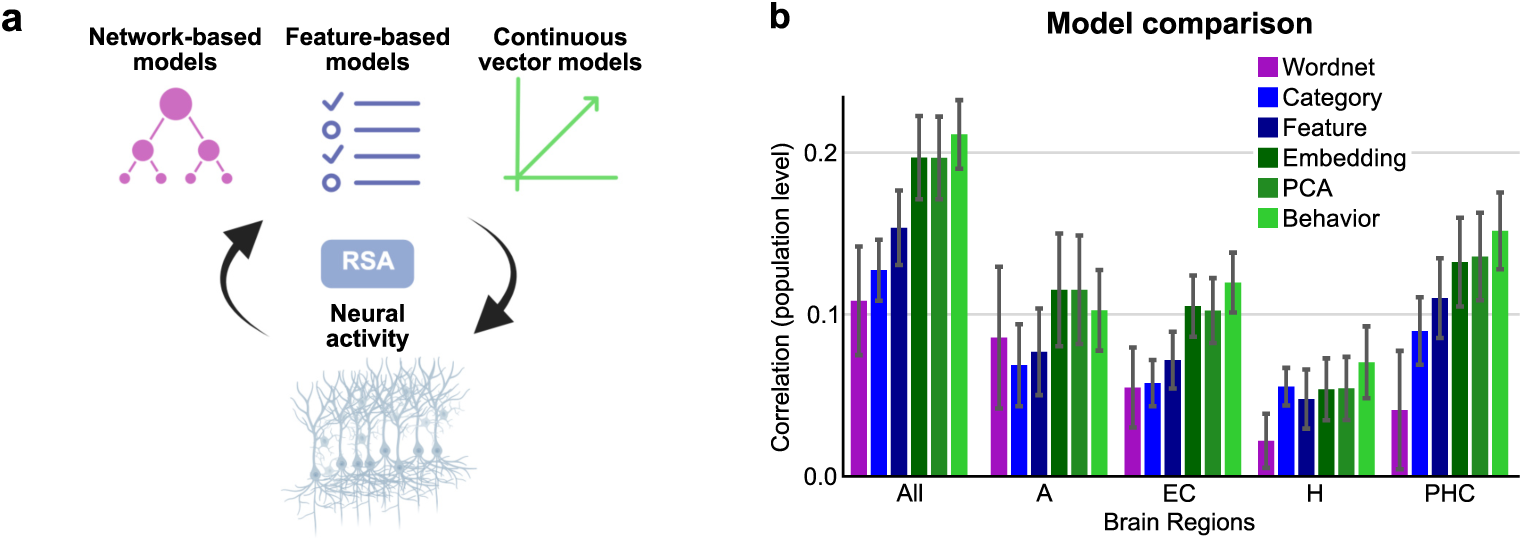
Predictive performance of different semantic models. a) An illustration of various theoretical approaches to semantic similarity, which are evaluated with respect to their performance in predicting neural activity. *Created in BioRender. Kehl, M.S. (2025)* https://BioRender.com/gvb6553. b) Spearman correlation between distances in semantic space and distances in neural population activation. The correlations are calculated for different distance metrics and brain regions. Error bars denote the bootstrap standard error.

To compare how well the three theoretical approaches predicted population spiking activity, we conducted a representational similarity analysis (RSA)^32^, which allows us to directly compare the distributed patterns of brain activity with semantic models at the level of representational similarities. To this end, we calculated the pairwise cosine distances between the neural activation vectors to all stimuli and correlated them with the pairwise cosine distances of the concepts in the respective semantic space (Fig. 4a). As this analysis requires a consistent set of stimuli (e.g., Mormann et al. ^15^) and our adaptive paradigm selects varying stimuli at later stages during one experiment session, we limited the population-level analysis to the 120 stimuli chosen for the initial experiment phase, which was identical for all patients.

For the network-based model, we chose *WordNet* ^33^ which provides a directed acyclic graph that captures semantic relationships between concepts and the Leacock-Chodorow Similarity as a metric for semantic relatedness. For the feature-based models, we made use of the feature and category descriptions provided by the THINGS dataset ^26^. The continuous vector embeddings were also taken from the THINGS dataset ^21^. These vectors are in a 300-dimensional space, but we also included PCA-reduced embeddings for comparison as they were part of our previous analyses (see *Methods*). Furthermore, we also tested a novel embedding in the THINGS database, derived from a behavioral odd-one-out-task performed by human subjects (see *Methods*).

Fig. 4b compares the ability of the three different theories to predict population spiking activity. Our analysis revealed that continuous vectors achieved the highest correlation with neural responses, while the network-based approach using WordNet performed the worst. Furthermore, the vector embeddings derived from human behavior (light-green) outperformed standard word embeddings derived from co-occurence statistics. When comparing between brain regions, it is noteworthy that the network approach performed comparatively better for the amygdala and poorly for the parahippocampal cortex, respectively. In addition, all models exhibited a noticeably low performance in the hippocampus compared to the other brain regions. Here, however, unlike in the other brain regions, discrete feature-based models performed comparably to continuous vector embedding models. Interestingly, the correlation over all brain regions was higher for all models compared to any single region.

In conclusion, this investigation into how different semantic theories account for MTL concept encoding reveals superior performance of continuous vector models of semantics. These models, which represent concepts as high-dimensional vectors, demonstrated the highest correlation with population-level neural activity. This suggests that semantic organization in the MTL aligns more closely with the distributed representations captured by these models, particularly those derived from human behavior. In contrast, network-based models performed least effectively, indicating a less prominent role for hierarchical graph-like structures in predicting neural responses.

### Multi-Field Semantic Tuning in Some MTL Neurons

The examples discussed thus far illustrated neurons tuned to a single semantic field. Interestingly, we also observed neurons that exhibited responses to multiple distinct concepts spanning different semantic categories. For instance, one neuron showed tuning for ”luggage” in a general sense, but also responded to certain electronic devices (Fig. 5a).

**Fig. 5:**
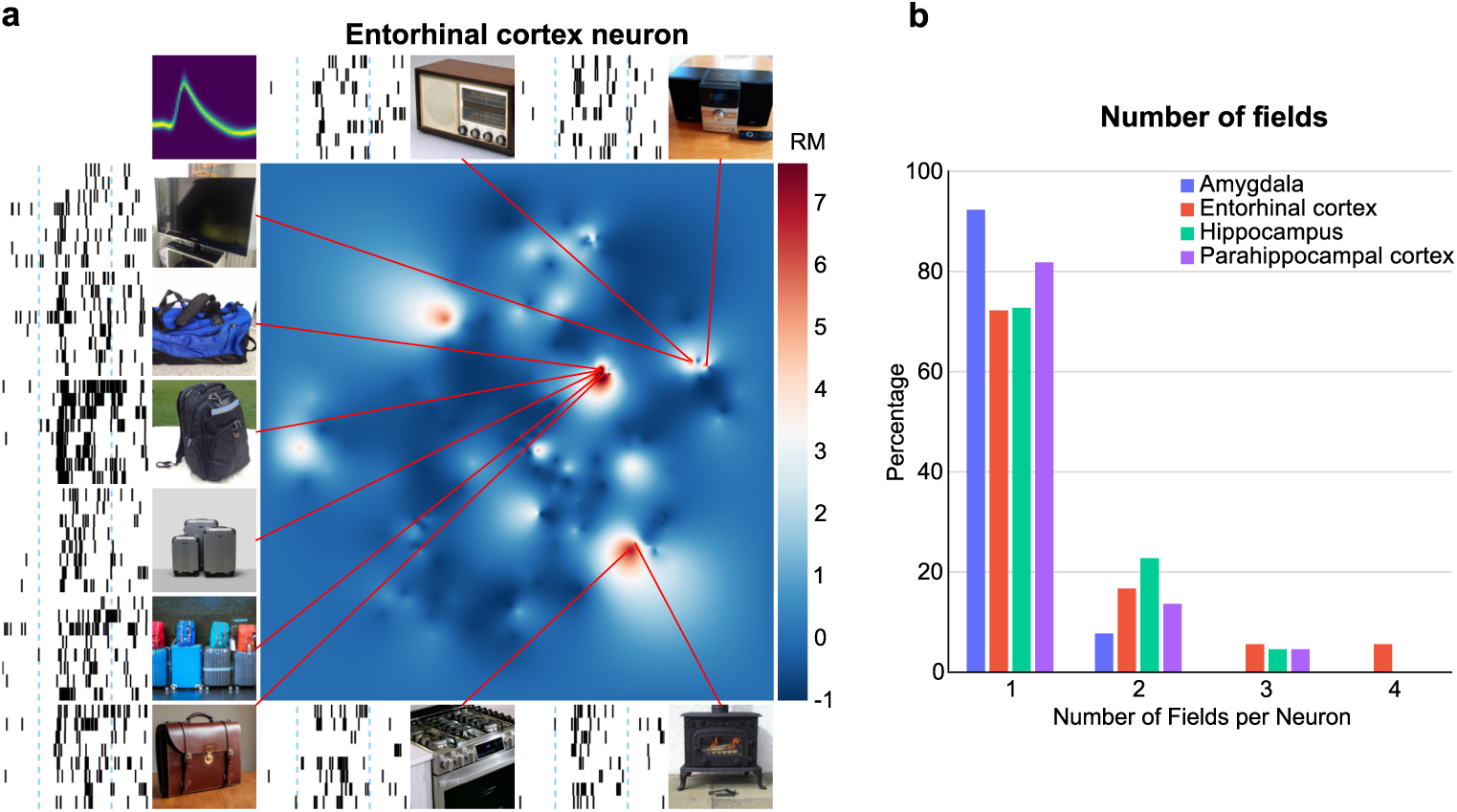
MTL neurons are occasionally tuned to multiple semantic fields. a) Tuning of example neuron #249 (entorhinal cortex). b) Percentage of neurons having different counts of semantic receptive fields for different MTL regions.

To systematically identify and quantify these multi-field responses, we analyzed the activation profiles of each neuron. We did so by counting the number of distinct peaks in the activation profiles of the neuron (*Methods*). A peak was identified as a significant increase in firing rate relative to baseline, indicating a response to a specific semantic cluster.

Fig. 5b summarizes the overall distribution of MTL neurons with one or more receptive fields. It shows that among those neurons exhibiting at least one defined semantic field, only a small percentage displayed two such fields, and very few neurons demonstrated three or more. This pattern of predominantly single-field tuning, with a minority of neurons showing two fields, was consistent across all investigated brain regions within the MTL (*χ*^2^(3, N=77) = 2.7, p = 0.44, testing for 1 field vs. more than 1 field). Specifically, the proportion of neurons with more than one field did not differ significantly between the hippocampus and entorhinal cortex (p = 1.00, Fisher’s exact test).

## Discussion

A fundamental question in cognitive neuroscience is how the human brain encodes abstract conceptual information to support complex human behavior. Concept cells in the human medial temporal lobe are thought to play a pivotal role in higher-level cognition, particularly in the processing of declarative knowledge and the formation, consolidation, and retrieval of episodic memories. However, the fundamental coding principles that govern their tuning to semantic information have remained largely unresolved. Understanding this tuning is crucial, as it provides a unique window into the neural basis of semantic representation at the single-cell level.

Given the constraints of limited patient time, the unknown semantic dimensions of concept-cell tuning, and the resulting infeasibility of using a fixed predefined stimulus set, we implemented a closed-loop experimental paradigm that adaptively selects stimuli by integrating real-time spike-sorting results. In this setup, we presented visual stimuli while recording single-neuron activity and analyzing spiking responses. Based on this online analysis, subsequent stimuli were selected from a high-dimensional semantic space to efficiently probe the boundaries and characteristics of a neuron’s semantic receptive field. This closed-loop methodology enabled a more targeted and detailed exploration of neuronal tuning compared to traditional paradigms that used preselected stimulus sets.

Critically, we found that MTL concept cells exhibit graded response profiles that scale with the semantic distance to their activation peak, revealing continuous semantic encoding at the single-neuron level. Our results indicate that concept cells exhibit narrow, graded tuning curves centered on specific regions of semantic space (receptive fields) with sharply reduced response activity to concepts farther from their tuning peak.

Since the earliest studies of sensory systems, tuning curves have been the standard tool for characterizing the contribution of individual neurons to population encoding by measuring their responses to relevant stimuli. Neurons in columns in the visual cortex were shown to be tuned to line orientation ^34,35^; auditory neurons to particular sound frequencies ^36–38^; motor cortex neurons to preferred directions of movement^39^; and, beyond sensory and motor systems, place cells in the rodent hippocampus responding to spatial locations ^40,41^. Our findings therefore extend knowledge about neuronal tuning to a new, specifically human, system where semantic processing and memory formation take place. Tuning curves may offer advantages over binary coding, potentially allowing neurons to convey richer, more nuanced information about a continuous range of semantic content in a more efficient way. Such graded representations would enable a smaller population of neurons to encode semantic information by leveraging population coding, where the combined activity of many neurons could more precisely represent a concept’s meaning. To achieve a similar precision, binary encoding would need a larger number of neurons, each responsive to only a specific concept. Furthermore, such a selective, but distributed coding may provide superior robustness to noise and graceful degradation in semantic processing, as information might not be reliant on the firing of specific units.

The utility of semantic tuning curves is particularly evident in their capacity for finer discrimination between closely related concepts and broader generalization across similar concepts. For discrimination, much like sensory acuity relies on the steepest slopes of tuning curves, neurons with steep semantic tuning curves allow subtle differences in semantic content to yield marked changes in firing rate, allowing the brain to distinguish between highly similar meanings (e.g., ”cup” vs. ”mug”; in contrast to the peak of the tuning curve, which might be a less sensitive region for fine discrimination given that its slope at the peak is zero; although both the peak and steepness have been suggested to be relevant in different conditions ^42^).

For generalization, a neuron’s response across a range of semantically related stimuli (e.g., various types of chairs) allows it to efficiently represent a category without needing a dedicated neuron for every specific instance. In essence, each neuron represents a ”consequential zone” ^43^ in semantic space, more or less wide, that may represent a hypothesis about which semantic cluster is currently relevant^44^. Such a broad coverage of semantic space at multiple scales would allow the brain to process novel, yet semantically similar inputs effectively, potentially allowing for a more flexible and adaptive understanding of the continuous complexity of the world’s meanings. Such a coding scheme could add more flexibility to the partially overlapping assemblies of binary neurons suggested by previous research ^45^.

If this distinction between discrimination and generalization is correct, then the steeper tuning curves in the hippocampus observed in our study might suggest a potential role for these high-slope regions in semantic discrimination, particularly if semantic inference operates under noisy conditions where subtle differences in input require clear differentiation. This, however, remains a point for further exploration.

Contrary to the graded tuning observed in our data, earlier work has suggested a binary, all-or-none coding. Rey et al. ^17^ reported a near-binary firing pattern: most of the normalized firing rates per neuron fell either onto 0 or onto 1 across stimuli, with only rare instances of firing rates in between (Fig. A.5, left panel). To reconcile the two findings, we reproduced their exact procedure with our data. We found the same overall pattern, but with a tendency for more ”in-between” firing rates and a lower percentage of response-eliciting stimuli in total (Fig. A.5, middle). The latter can be explained by the fact that the authors presented stimuli which had already been identified as response-eliciting in a preceding screening experiment. However, due to the normalization of the firing rates there will always be, by definition, a single stimulus with a response strength of 1 per responsive neuron. This methodological artifact is amplified by the sparse sampling of the underlying concept space – in the absence of a very dense sampling of concepts, or of a closed-loop adaptive procedure similar to ours, experiments are likely to conclude that many neurons only respond to a single stimulus with a maximal response magnitude. To test how a neuron responds to other stimuli apart from the one used for normalization (whose response is 1 by definition), we re-analyzed our data after excluding the respective stimulus for each responsive neuron (Fig. A.5-Right). The result revealed a broad distribution of firing rates, without a preference for the firing rate of 1. This finding challenges the notion of strict binary coding in the human MTL. Previous suggestions of binary behavior, we argue, are better explained by an artifact of normalization combined with sparse stimulus sampling within the semantic space.

Investigating responses across different MTL regions revealed notable variations in semantic tuning and representation. Specifically, the amygdala showed sensitivity to the category of animals ^15^ (Fig. 3b), though its representation was less pronounced compared to food. In contrast, the parahippocampal cortex showed the highest sensitivity to the categories of animals and vehicles, in line with the region’s known involvement in processing scene information ^16^. While not central to the current study, future research should disentangle effects of category and background to more clearly identify regional selectivity.

Our results also shed light on the long-standing debate about how concepts are represented in the human mind. We contrasted three different approaches to lexical semantics: network-based models, feature-based models, and continuous vector models based on distributional semantics and behavioral data. The results revealed a superior fit of continuous vector embeddings over network and feature-based models. The data therefore supports the relevance of continuous vectors rather than strict hierarchies. Note, that the best fit was obtained by a comparably low-dimensional vector space directly derived from human behavioral judgments (see also Fig. A.7) without any assumption that the brain compiles statistics based on word co-ocurrence. Overall, the strongest conclusion from our research is that vector space models, with individual neurons showing continuous responses to specific vectors, provide the best fit to human concept cells. This conclusion fits with research on large language models (LLMs), where concept organization within high-dimensional embeddings reflects distributional semantics^46,47^. While some studies explore potential hierarchical structures in LLM representations, e.g., Park et al. ^48^, these were restricted to small portions of the semantic space.

The idea that neurons encode vector spaces has gained attraction in recent experimental and theoretical work^49–51^. In particular, an extensive analysis of the tuning of face-responsive cells in macaque monkeys has also suggested that their firing profile is best described by an underlying high-dimensional vector space for face shape, in this case with 50 dimensions ^52^. However, there is an interesting potential difference: our findings in MTL cells point towards a ”peak” coding scheme where neuronal firing reaches a peak at a preferred point in space (e.g., Fig. 2a&b, Fig. A.6), whereas face-responsive neurons seem to support an ”axis” model whereby neuronal response increases monotonically along a distinct direction in vector space. We note, however, that what looks like a local tuning peak when the stimulus space is depicted in two dimensions (as in Figs.2a&b) may actually correspond to a single axis in the full underlying high-dimensional space. The distinction is made more difficult by the fact that in semantic space, unlike face space, not all vectors correspond to actual concepts. For instance, it seems impossible to present a picture that would be ”more fiery” than the fire-related images in Fig. 2b – whereas it is always possible to distort a face into a caricature that corresponds to a larger face vector see, e.g., Leopold et al.^53^. Thus, a comprehensive comparison and disambiguation of peak versus axis coding strategies will necessitate dedicated future experiments.

While the vast majority of concept cells analyzed displayed a single, well-defined semantic receptive field, we also encountered rare instances of neurons responding strongly to multiple, distinct regions within the semantic space. Although this observation might suggest speculative parallels with spatial coding schemes like place and grid cells, we highlight several points of caution: such multi-field responses are rare in our dataset, they did not exhibit a fixed geometric structure, and we found no difference in the percentage of multi-field responses across different MTL regions (in particular, between the hippocampus and entorhinal cortex).

In our study, each concept was presented multiple times using diverse images. The fact that human MTL neurons fire to such diverse pictures, as long as they fall within a local region of semantic space, fits with previous work on concept cells demonstrating abstract responses to both words and pictures ^8,9,17,54,55^. This suggests a tuning to the semantic content of objects rather than purely visual features, as suggested by recent studies ^19^. A limitation of our study, however, is that we did not explicitly decorrelate visual features from semantic content. Still, the observed invariance in responses across varied visual instantiations of the same concept provides strong indications of semantic coding. Moreover, the identified similarity in responses to different objects with strong semantic relationships (e.g., onions and cakes), despite their visually distinct features, further supports that these neurons encode abstract information. Due to constraints on the duration of the experiment, we did not investigate whether these neurons also respond to object names or stimuli from other modalities. Future studies could more directly test the distinction between visual and semantic coding by employing stimuli designed to isolate visual features from semantic content, and additionally explore responses to multimodal abstract representations.

Finally, a critical consideration in our study is the underlying geometry of the semantic space. The choice of which semantically related stimuli to present next, in our paradigm, was based on word embeddings that were matched to the THINGS dataset. This choice inherently assumes a specific organization of semantic information. For future studies, it will be important to probe whether different choices affect the results. In conclusion, our study provides a detailed characterization of semantic tuning in MTL concept cells, revealing graded response profiles with regional variations within the MTL. These findings thus advance our understanding of how abstract knowledge is neurally encoded. Furthermore, the novel closed-loop paradigm developed here offers a powerful tool for future investigations into the fine-grained receptive fields of concept neurons. The framework we created can be extended to other types of stimuli and sensory modalities, opening the door to exploring concept-neuron behavior more comprehensively and across varied dimensions of conceptual space. This has broad implications for understanding the fundamental principles of semantic representation in the human brain.

## Methods and Materials

### Ethics Statement and Participants

The study was approved by the Medical Institutional Review Board of the University of Bonn (accession number 095/10 for single-unit recordings in humans and 287/20 for the current paradigm in particular) and adhered to the guidelines of the Declaration of Helsinki. Each patient gave written informed consent.

We recorded during 36 sessions from 15 neurosurgical patients (5 females, 10 males; ages 22-60, average age 39.3) implanted with depth electrodes for pre-surgical invasive seizure monitoring. Each experimental session lasted approximately 40 minutes (mean: 39 minutes, SD: 15 minutes).

### Electrophysiological Recordings and Spike Sorting

Nine microwires (8 high-impedance recording electrodes, 1 low-impedance reference; AdTech, Racine, WI) protruding from the tip of the depth electrodes by approximately 4mm were used to record signals from MTL neurons. The total number of implanted microwires per patient was on average 95.33 (SD = 10.73) and ranged from 72 to 104. Out of these, an average of 73.33 (SD 7.89, minimum=64, maximum=80) microwires per patient were located in the MTL target regions and included in this study. Signals were amplified and recorded using a Neuralynx ATLAS system (Bozeman, MT), at a sampling rate of 32,768 Hz, and signals were referenced against one of the low-impedance reference microwires. Spikes were first automatically extracted and sorted using the Combinato software (https://github.com/jniediek/combinato) ^56^. The result was manually verified and clusters labeled as artifacts, single- or multi-units using the Combinato GUI. The *Duplicate Event Removal algorithm* ^57^ was applied to all sessions after manual spike sorting.

### Stimuli and Embeddings

Subjects were presented with visual stimuli from the THINGS dataset, which provides semantic data for 1,854 concepts named by participants during extensive online surveys ^20^. For each concept, a set of at least 12 images in a natural context is provided as well as the corresponding synset from WordNet, 300 dimensional semantic vector embeddings based on word2vec refined with knowledge about different senses based on WordNet (if the corresponding concepts could be identified) ^21^. In addition, 27 categories were assigned to the concepts. It should be noted that a concept can have several categories, e.g., ’aquarium’ which is both labeled as ’container’ and ’home decor’, or no category. Each concept is also attributed with descriptive features such as ’is made of metal’, ’has wings’, or ’is round’ ^26^. The total number of distinct features for the concepts in the THINGS dataset is 9,508.

We used the THINGS embeddings ^21^ and reduced the number of dimensions using principal component analysis (PCA) from 300 to 25. We used the cosine distance as a high-dimensional distance metric, normalized the result with the smallest and largest distance between all concepts, and calculated 1 minus this distance as a metric for semantic similarity. In a separate, but related, project, the THINGS initiative added behavior-related embeddings to the dataset, which were generated with human subjects performing an odd-one-out task ^31^. We used the resulting 66-dimensional embeddings for the model comparison in Fig. 4i.

### Paradigm with Online Spike Sorting

To explore the semantic field of a single neuron, we developed an experimental setup which allows for dynamic stimulus selection during the experiment by performing real-time data analysis. To this end, the neural signals recorded on the recording PC were additionally transferred to the experiment laptop used by the patients, where the algorithm analyzed them and determined which stimuli to present next. For a similar setup previously developed in our lab, see Knieling et al. ^58^.

A trial entailed the presentation of a blank screen for a random duration in the range of 200ms – 400ms, then a fixation dot in the center of the screen for 300ms, followed by the image that remained on screen until the subject responded with a button press. To ensure engagement, participants performed a simple task: indicating with left/right arrow presses whether an object was liftable (e.g., a bottle) or not (e.g., a tree). This kept their focus on the images, yet they remained oblivious to the intricate, neural-activity-driven closed-loop system operating in the background to select the next stimuli.

The experiment was structured into three phases:

***Phase 1: Exploration***

Initially, the subject was presented with a set of stimuli providing a large coverage of semantic space. These stimuli were selected by applying k-means clustering with k=120 to the full 300-dimensional embeddings provided by the THINGS database ^21^, and selecting the stimulus closest to the center of each cluster to cover the semantic space. We started the experiment with 150 initial stimuli taken from another paradigm used in our lab and switched to the 120 initial stimuli chosen with this approach after the first three patients in order to leave more time for the exploration of semantic space. Each initial stimulus was shown between 4 to 6 times, thus yielding a first estimate of which stimuli do not elicit a response in any of the recorded neurons and can be eliminated.

***Phase 2: Response Consolidation***

The most promising stimuli were chosen and presented to the patient again in order to attain a total of 8 trials for each of these stimuli.

***Phase 3: Local Search***

The strongest response-eliciting stimuli were selected, and concepts in their semantic vicinity were presented to the patient as illustrated in Fig. 1e. This last step was repeated iteratively in order to discard stimuli for which no response-eliciting neighbors could be found and/or to replace the starting points of the semantic search with even better stimuli. During each step of the last phase, the 4 to 8 most promising stimuli were chosen, unless the respective neuron failed to respond to the last 2 presented stimuli in the searched neighborhood. We selected the most promising stimuli based on the lowest alpha value for any stimulus in the binwise ranksum test (see below) used to determine responsiveness of neurons. For each of the picked stimuli, their closest neighbor was presented to the patient.

When we started recording data with this setup, we used a relatively complex algorithm to determine the next stimuli (instead of using nearest neighbors). Our initial goal was to look for a local maximum, i.e., the stimulus evoking the highest response in the neuron and we did not know how far the tuning width of a neuron extends. Thus, we systematically probed the neighborhood of the initial stimulus at varying distances until a stimulus evoking a stronger response was found and was then used to replace the initial stimulus. However, it quickly became apparent that the responsive fields of a neuron are commonly confined to a rather limited area, such that presenting stimuli at higher distances proved unhelpful. Therefore, we switched to using the nearest neighbor approach (see above) after the first five patients.

During the last iteration of the Phase 1, the Combinato software was used to analyze the neural data acquired so far, automatically identifying putative neurons and assigning them the recorded spikes. For the data recorded in later stages, template matching was used to assign the spikes to cells. All other analyses to calculate the responses were executed in real-time using CUDA in order to substantially accelerate the process with the help of graphics processing units (GPUs).

The experimental setup included adjustable parameters that allowed the session length to be tailored to each participant’s level of comfort and capacity for engaging with and focusing on the task. The parameters which could be adjusted were:

- The number of initial trials during the first phase
- The number of stimuli selected for the second phase
- The number of iterations during the third phase
- The number of stimuli presented in each iteration of the third phase

### Analyses and Stimulus-Delivery Software

The stimuli were presented to the patient using psychtoolbox3 (www.psychthoolbox.org) and Octave 5.2.0 (www.gnu.org/octave) on a Windows 10 system on a high-end Laptop. Combinato was used for online spike sorting. The program used for response calculation and stimulus selection was implemented in C++ with VisualStudio, additionally using the NVIDIA CUDA toolkit to increase computing power. A Python script served as an interface between the C++ program and Combinato. The offline analysis was performed in MATLAB and Python3 using a DataJoint database system ^59^.

### Statistical tests

Unless stated otherwise, all error bars denote the mean and standard error of the mean (SEM). Significance tests were executed after an *analysis of variance* (ANOVA) test showed significance (p-value *<* 0.05) and were done as pairwise t-tests. No significance is defined as a p-value *>* 0.05. A single asterisk (*) denotes 0.01 *<* p-value ≤ 0.05, double asterisks (**) indicate 0.001 *<* p-value ≤ 0.01, and triple asterisks (***) represent p-values ≤ 0.001.

### Neuronal responses

Neurons were classified as responsive based on established criteria ^14,60^. Specifically, we defined a neuron to be responsive to a stimulus based on a binwise rank-sum test with additional correction for multiple comparisons using the Simes procedure as previously described ^60^ (with *α <* 0.001). We used the 1000ms after stimulus onset as relevant time window to calculate responsiveness and firing rates. The firing rate of a neuron is the number of spikes per second in the response time window, averaged over all trials a stimulus was presented in. Additional criteria for a neuron to be considered responsive to a stimulus were at least one spike being present in at least 50% of stimulus trials, and an average firing rate above baseline and ≥1Hz in the response time window.

Neurons which were responsive to at least one concept (presented through different pictures of that concept) were characterized as responsive and included in the analysis. Normalization of firing rates was done using the minimum and maximum firing rates of a neuron to any stimulus, so that the normalized firing rate always has a value between 0 and 1. The response magnitude of a neuron to a stimulus is expressed by a z-score using the mean and standard deviation in the baseline interval.

### Tuning

#### Peak and Semantic Similarity to Peak

We calculated an estimated peak in embedding space by calculating a weighted average of the three stimuli evoking the highest firing rate; with the firing rate serving as respective weight per stimulus. The semantic similarity to the peak was then 1 - cosine distance of each stimulus to the estimated peak with the cosine distance normalized with respect to the highest and lowest cosine distance between any two embeddings of the dataset.

#### Neural Activation Heatmap

The 2D activation heatmaps created for each neuron were based on the t-SNE reduced space from the THINGS database ^20^. The color-coded 2D map was created using linear interpolation of the z-scored response magnitude for each stimulus.

#### Single Neuron Tuning Fit

For each responsive neuron, we fit a logistic sigmoid function to the normalized firing rate as a function of the semantic similarity to the peak using scipy’s curve fit package and the function *expit*(*k* ∗ (*x* − *x*0)) with the bounds 0 ≤ *x* ≤ 1 and −100 ≤ *k* ≤ 100. We uses *x* = 0.5 and *k* = 1 as initial values. The tuning width was calculated as 1 − *x*0, and the tuning steepness defined as the slope at the point of inflection, which characterizes the steepest slope of the fit.

#### Coactivation probability

For the coactivation probability, we started from each response-eliciting stimulus and, from there, split the remaining stimuli by their semantic similarity into fixed intervals [0 : 0.1 : 1.0]. The coactivation probability was then defined by the counts how often a stimulus with a certain similarity to a response evoked a response, divided by the counts how often a stimulus with this similarity was presented to a given neuron.

#### Split Spearman Correlation

For the Spearman correlation split by semantic similarity to the peak, we used the single neuron tuning split into intervals [0 : 0.1 : 1.0], and calculated the Spearman correlation between the normalized firing rate and the semantic similarity to the activation peak for each interval separately per neuron. We accumulated the values per interval across all neurons using the mean and SEM.

#### Linear Regression

For the linear regression models, we first used RidgeCV from Python’s scikit-learn package. To determine a suitable regularization size (the alpha parameter) for the model, we used nested cross validation, using a split of 80%*/*20% of stimuli for training and validation sets. We used the *R*^2^-value for model evaluation. To get the feature importance score for a specific feature, we used the leave-one-feature-out (LOFO-)score and re-trained the model with the same split in training and test sets removing said feature from the model. The resulting drop in *R*^2^ for removing a feature characterizes the LOFO-score. We created linear models for each neuron using the categories, features, and PCA-reduced embeddings described earlier.

#### Multi-Fields

To determine the number of fields a neuron responds to, we created 2D heatmaps interpolating the alpha values of all stimuli resulting from the rank-sum test defined earlier instead of the response magnitudes. We defined a responsive field as a connected area with p-value *<* 0.001 including at least two stimuli evoking a response in the neuron and counted the resulting fields per neuron.

## Data and code availability

Data and code to reproduce the main findings are available upon request.

## Acknowledgements

We thank all patients for their participation in this research. This research was supported by grants from the DFG (MO 930/4-2, MO 930/15-1, SFB 1089, SPP 2205, SPP 2411) and a NRW Network Grant (iBehave) (F.M.). Icons used in the figures were obtained from the Noun Project (https://thenounproject.com) under a NounPro subscription.

## Author contributions

K.K. and F.M. designed the study. K.K. implemented the study. R.S. and F.M. recruited patients. V.B. and F.M. implanted the electrodes. K.K. and Y.Q. performed the experiments. K.K. collected and analysed the data. K.K., S.D., Y.L., and F.M. planned and designed the analysis with contributions from M.S.K., P.K., M.N.H., and T.R.; K.K., P.K., and M.S.K. designed and prepared the figures; K.K. and Y.L. wrote the paper with contributions from M.S.K., A.D., P.K., M.N.H., T.R., S.D., and F.M. All of the authors discussed the results and commented on the manuscript.

---

**Fig. A.1:**
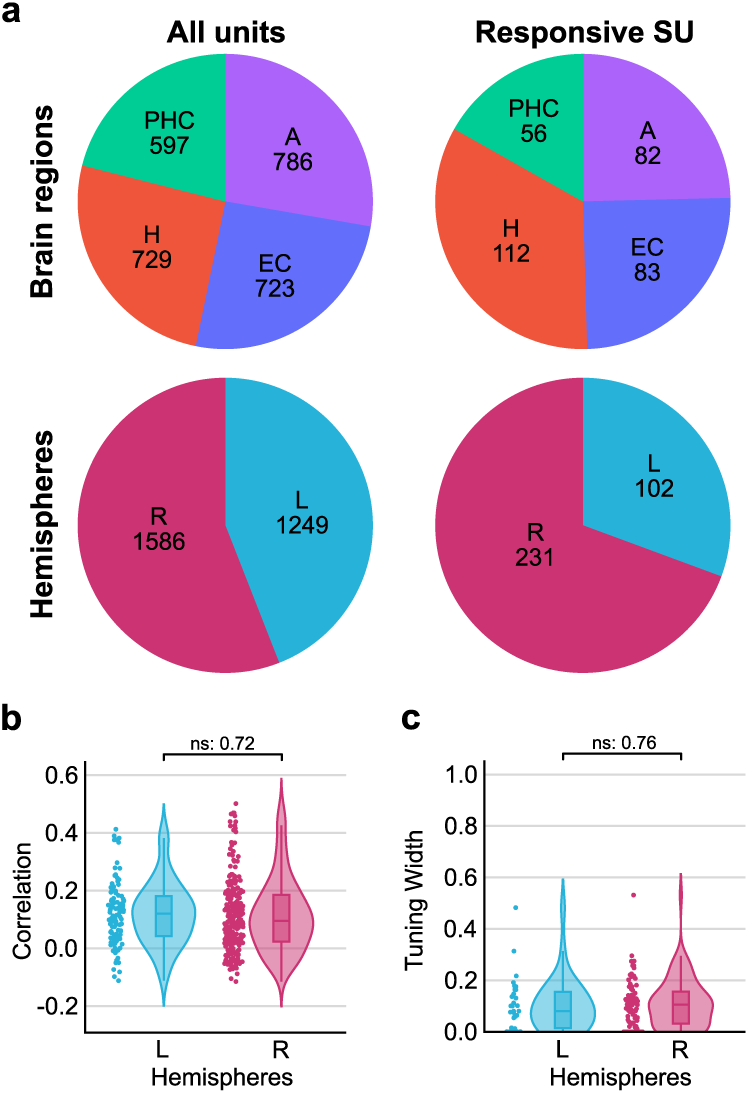
Additional single unit and tuning characteristics. a) Total number of recorded units and number of responsive single units by brain region and hemispheres. b) Distribution of Spearman correlation between semantic similarity to activation peak and response magnitude for all neurons by hemispheres. c) Tuning width for all neurons by hemispheres.

**Fig. A.2:**
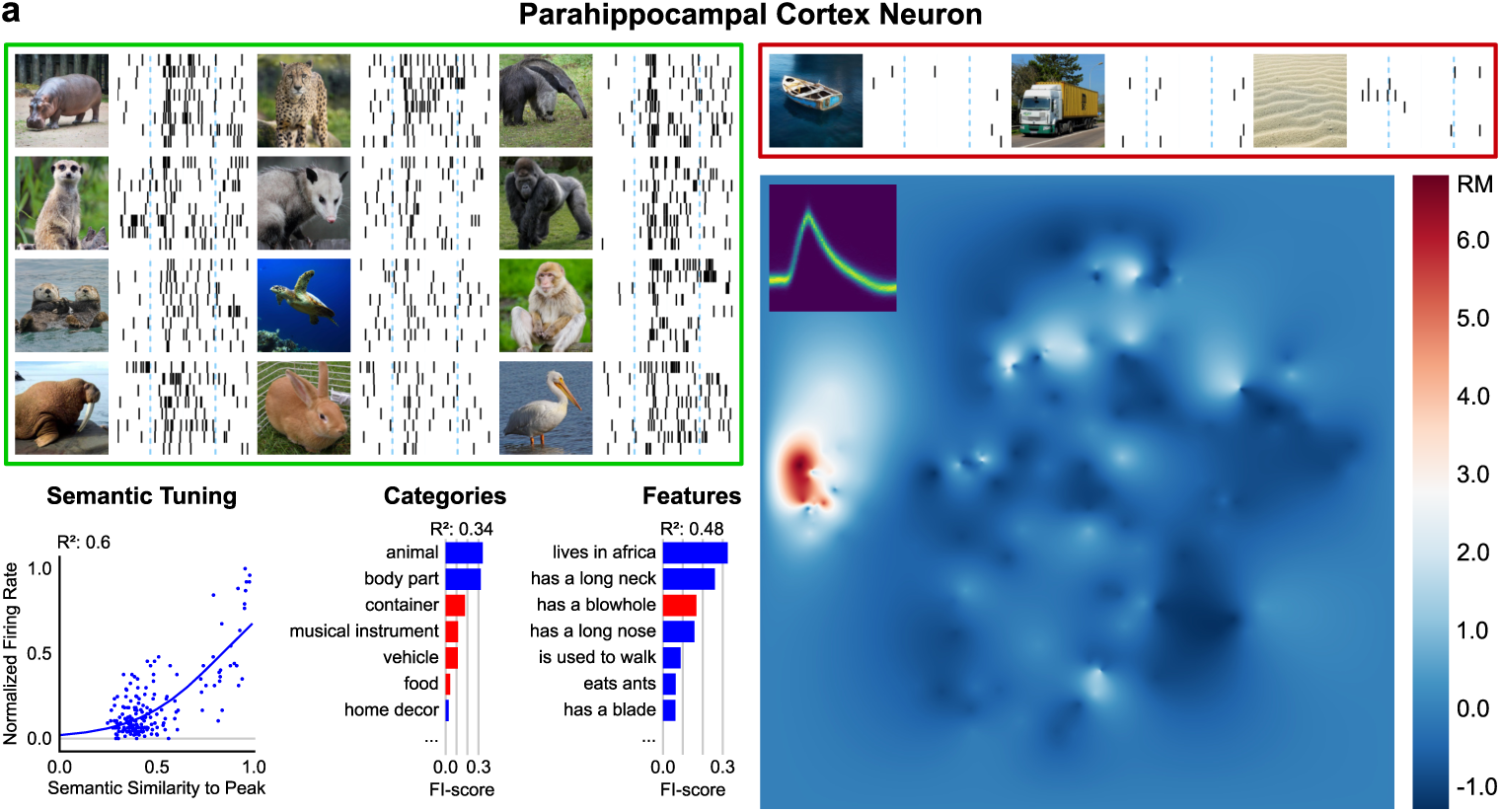
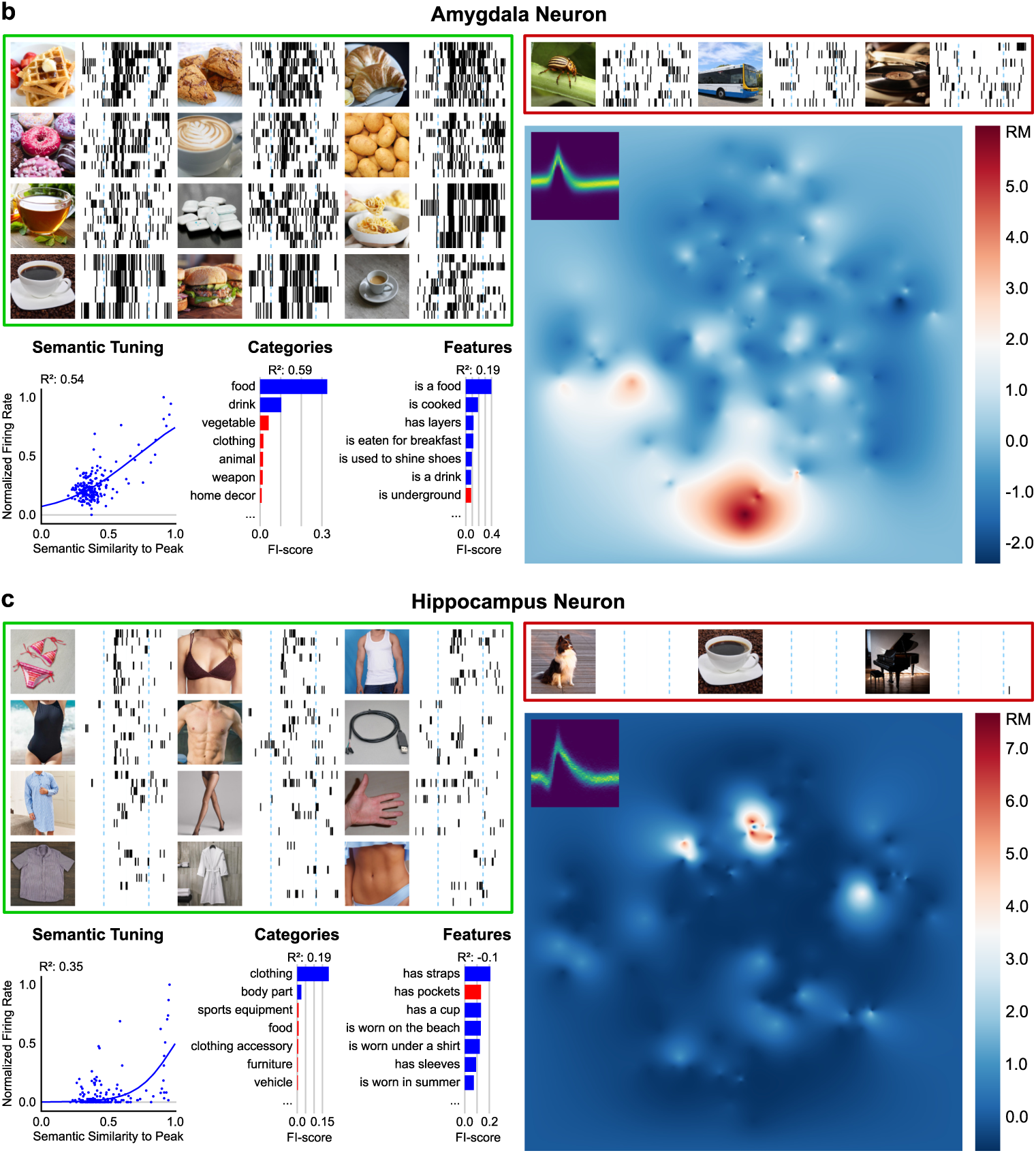
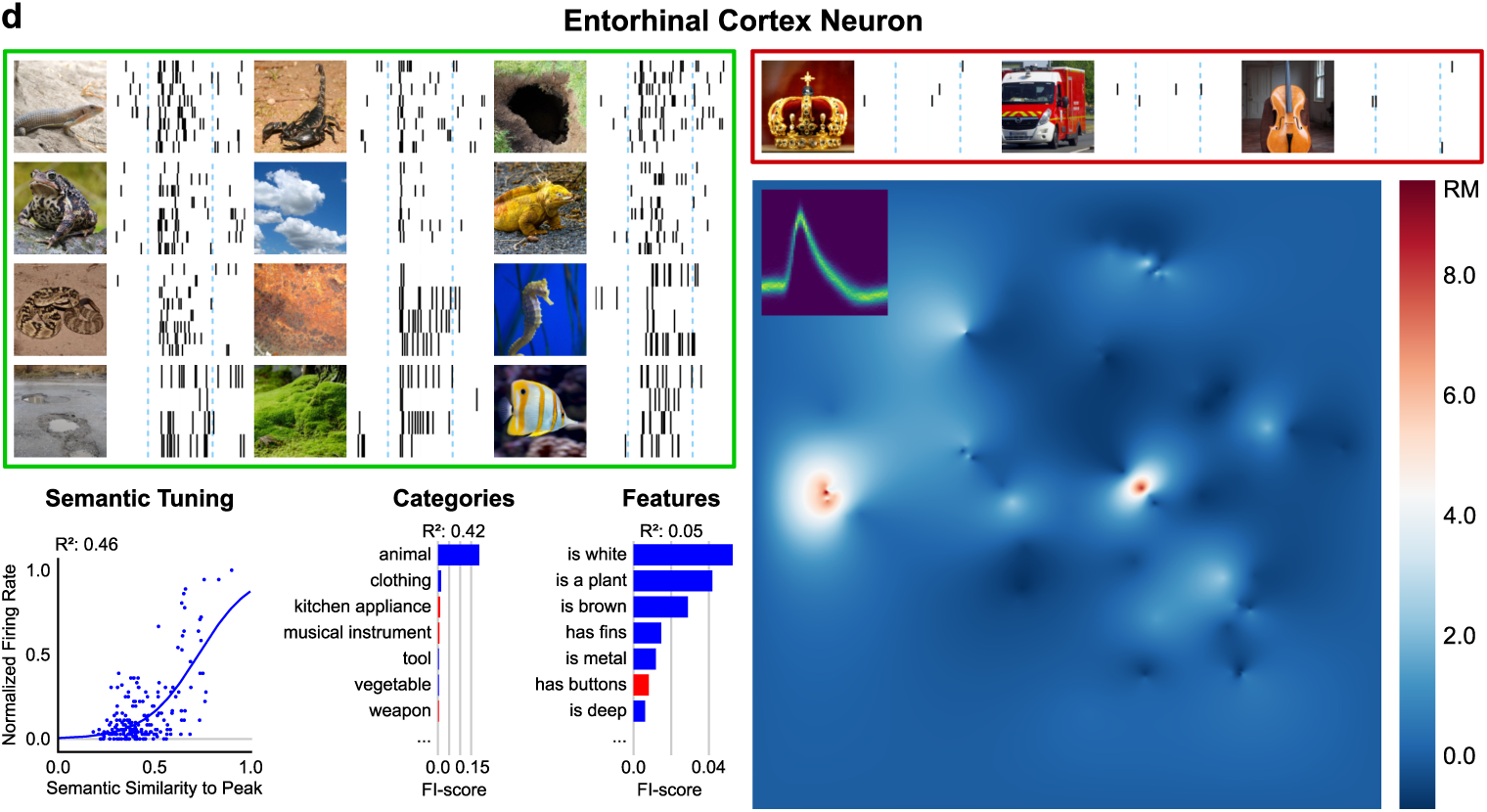
Response behavior of example neurons in the human MTL. a) Shows neuron #280 responding to animals (parahippocampal cortex) including spiking activity to several stimuli (green box: The 12 stimuli with lowest alpha-value in the binwise ranksum test; red box: 3 stimuli the neuron did not respond to), the semantic tuning curve (as in Figure 2a top right), FI-score for categories and features (as in Figure A.4), and the activation heatmap (as in Figure 2a left). b-d) Same as a, but for neurons #60 (amygdala), #254 (hippocampus) and #7 (entorhinal cortex).

**Fig. A.3:**
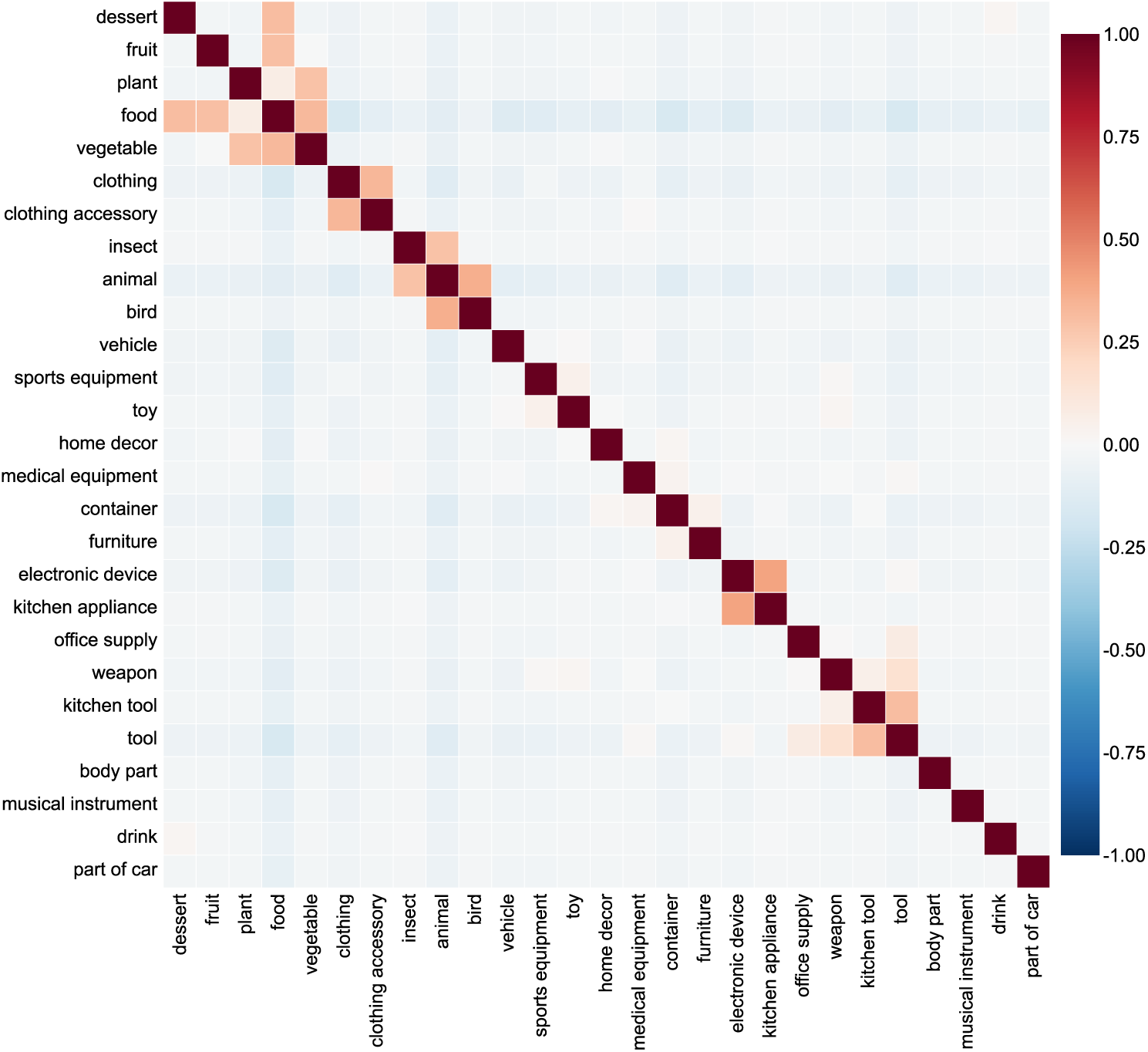
Correlation matrix between categories. The matrix shows Spearman correlations between categories, calculated from the concept–category matrix of the THINGS-dataset, which encodes whether each concept belongs to a given category.

**Fig. A.4:**
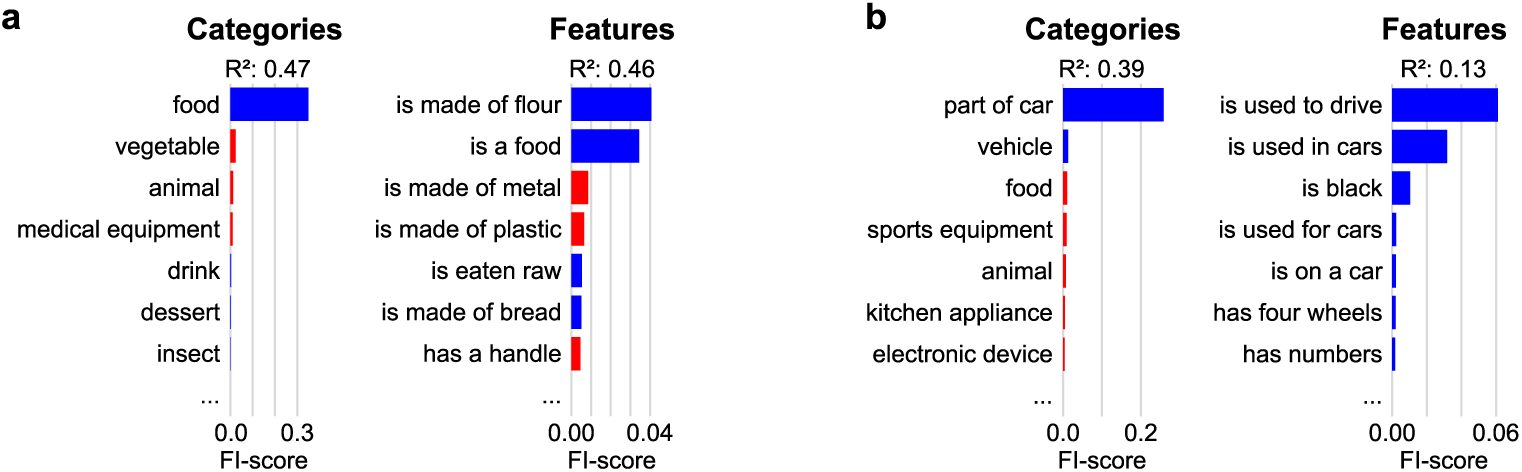
Exemplatory category FI-scores of single neurons in the human MTL. a) Top 10 categories (left) and features (right) with highest FI-score in a regression model for example neuron #75 presented in Figure 2a. Blue/red indicate positive/negative effects on firing rate. *R*^2^ represents the model’s performance in explaining unseen data. b) Same as a, but for example neuron #191 presented in Figure 2b.

**Fig. A.5:**
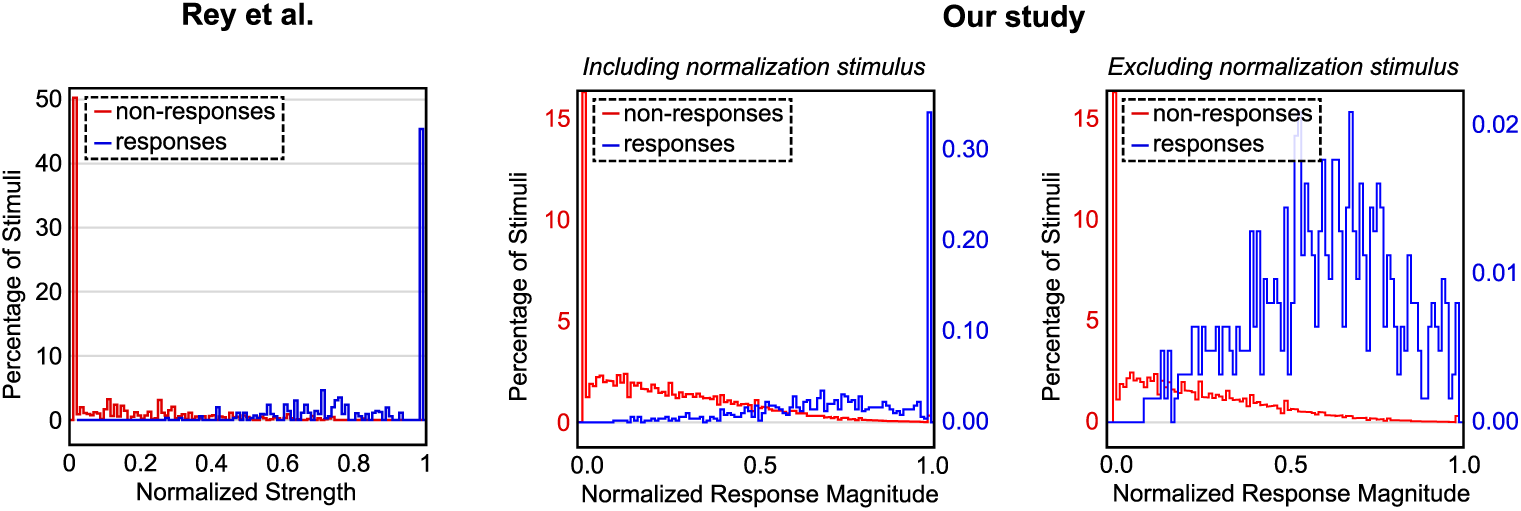
Binary vs graded responses in concept cells. The figures compare data from Rey et al. ^17^ and our own recorded data. They illustrate the normalized response magnitude for all responsive neurons split by stimuli based on whether they induced a response. The right figure excludes the normalization stimulus (peak stimulus) and shows that the remaining responses do not appear as a flat horizontal line.

**Fig. A.6:**
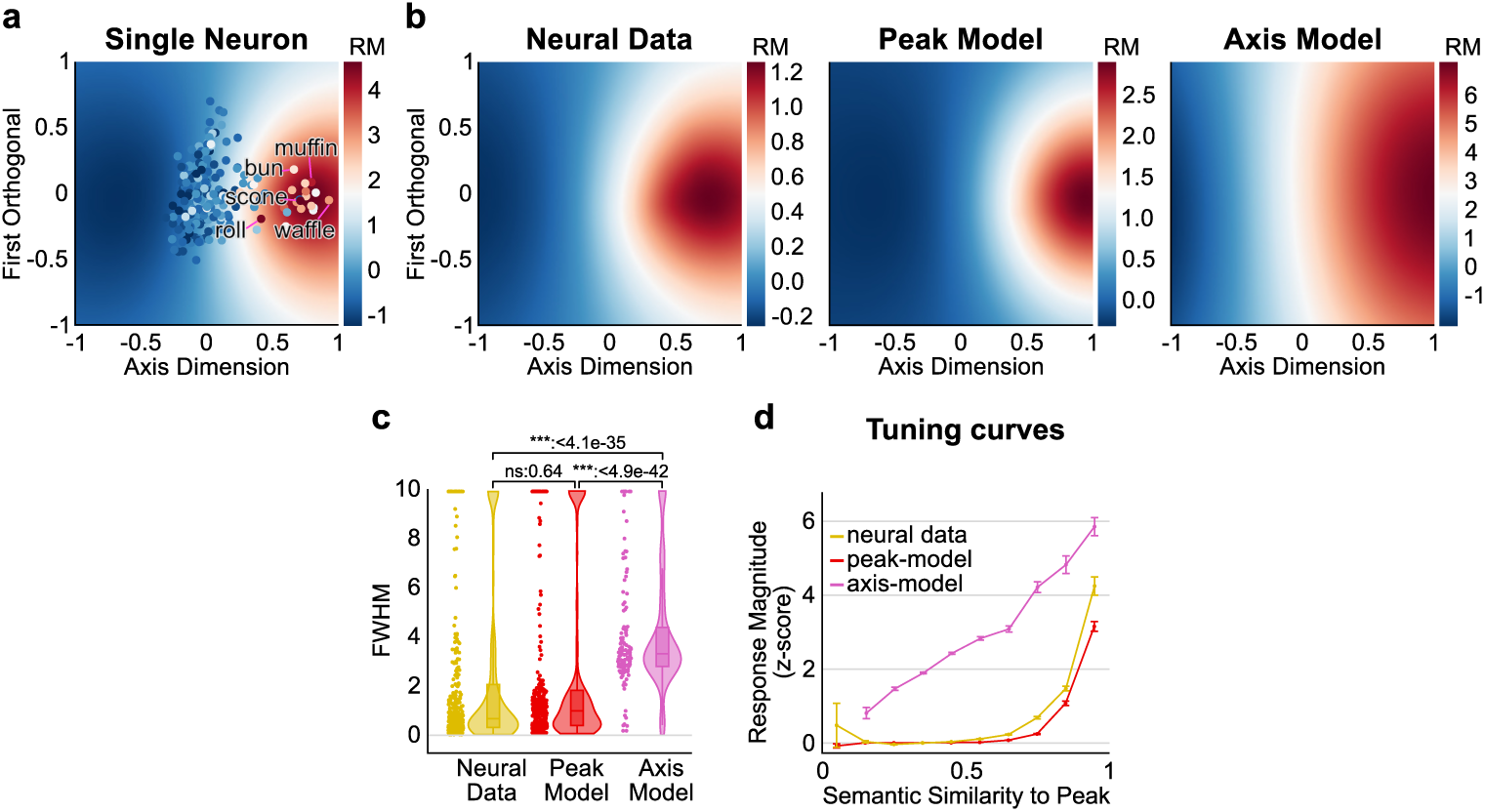
Comparison between simulated peak- and axis-neurons and real neural data. To compare whether the axis- or peak-model fits better to the recorded data, we simulated one neuron per recorded neuron following each model. For simulating the peak model, the response magnitude decreases according to a Gaussian function with distance to a peak embedding; for the axis model, the response magnitude increases along an axis fit with linear regression to the original’s neurons responses. a) We fit a Gaussian model and aligned the first axis of the PCA reduced embeddings with an optimal axis fit using linear regression to predict the neuron’s firing behavior. We visualized the first two dimensions of the prediction of the Gaussian fit (which are the dimension following the axis fit and the first orthogonal dimension) for the data recorded for neuron #75 presented in Figure 2a. Colors represent the predicted response magnitude (RM) of the model; the dots show the actual concepts presented to the neuron colored according to the actual response magnitude of the neuron. b) We averaged the resulting heatmaps described in a) for all neurons for the different models. The plot for the neural data shows a closer resemblance to the peak-than to the axis-model, as the average predicted response magnitude decreases along the first orthogonal from a center peak activation. c) To quantify this effect, we calculated the full width at half-maximum (FWHM) of the first orthogonal axis of the Gaussian fits through each individual peak. For visualization, values greater than 10 were truncated to 10. The neurons simulating a peak-model do not show a significant difference to the neural data in an independent t-test, while the axis-model shows a significantly higher FWHM on average. d) Shows the semantic tuning curves as in Figure 2d for the three models. The axis-model shows a rather linear increase in the response magnitude towards the peak embedding, while the slope is rather low for higher distances and high for lower distances to the peak embedding in the peak-model and the neural data.

**Fig. A.7:**
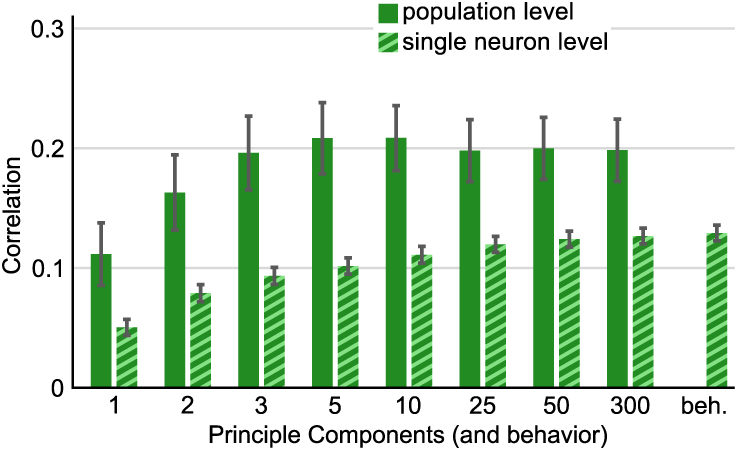
The influence of the dimensionality on the model’s performance. Given the strong performance of continuous vector embeddings (Fig. 4b), we also investigated how the dimensionality of the semantic space influences model performance and compared the spearman correlations for different counts of principal components given by PCA. The population-level analysis is calculated as in 4a. The single neuron level analysis shows the correlation between the response magnitude and the similarity to the activation peak for each neuron. On the right, we also included the result for the behavior-related embeddings ^31^. We found that even models with a relatively low number of dimensions (e.g., 5 or 10 principal components) performed comparably well or even better than the full-dimensional embeddings. However, our primary single-cell analyses focus on the correlation between semantic similarity to the activation peak and single-neuron activity. In this context, increasing the number of dimensions generally led to better results, with the behavioral embeddings again outperforming semantic embeddings (Fig. A.7, single-neuron level). While these two distinct outcomes mean there is no single ”optimal” number of dimensions, we chose 25 principal components as a compromise for representing the semantic space. Importantly, our results were robust and reproducible with other numbers of principal components and with the behavioral embeddings (see Fig. A.8).

**Fig. A.8:**
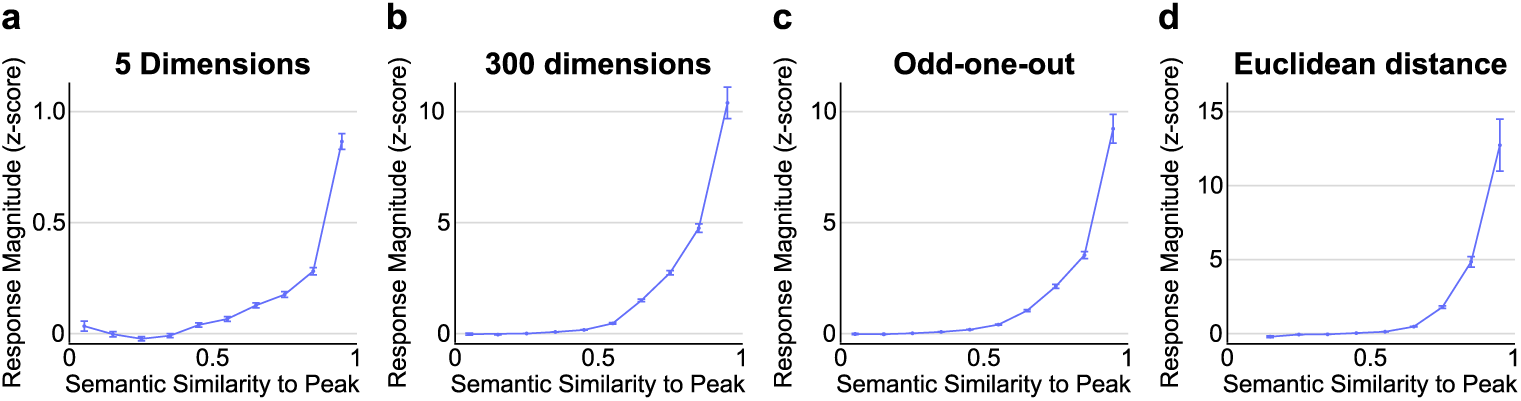
Tuning curves for alternative distance metrics. The tuning curves were calculated as in Figure 2d, but using alternative distance metrics to measure semantic relatedness of concepts. a) THINGS-embeddings were reduced to five dimensions using PCA. b) The full 300 dimensional THINGS-embeddings were used. c) Shows the behavior-driven embeddings created by an odd-one-out task^31^. d) THINGS-embeddings were reduced to 25 dimensions using PCA; with the euclidean instead of the cosine distance as metric for semantic similarity.

## References

[1] Posner, M. I. & Keele, S. W. On the genesis of abstract ideas. Journal of experimental psychology 77, 353 (1968).

[2] Medin, D. L. & Schaffer, M. M. Context theory of classification learning. Psychological review 85, 207 (1978).

[3] Murphy, G. The Big Book of Concepts (MIT Press, Cambridge, MA, 2002).

[4] Rosch, E. & Lloyd, B. B. Cognition and categorization Psychology Revivals Series (Routledge, Abingdon, Oxfordshire, UK, 2024).

[5] Squire, L. R. & Zola-Morgan, S. The medial temporal lobe memory system. Science 253, 1380–1386 (1991).

[6] Squire, L. R., Stark, C. E. & Clark, R. E. The medial temporal lobe. Annu. Rev. Neurosci. 27, 279–306 (2004).

[7] Quiroga, R. Q., Reddy, L., Kreiman, G., Koch, C. & Fried, I. Invariant visual representation by single neurons in the human brain. Nature 435, 1102–1107 (2005). Publisher: Nature Publishing Group.

[8] Quiroga, R. Q., Kraskov, A., Koch, C. & Fried, I. Explicit encoding of multimodal percepts by single neurons in the human brain. Current Biology 19, 1308–1313 (2009).

[9] Kehl, M. S. et al. Single-neuron representations of odours in the human brain. Nature 634, 626–634 (2024).

[10] Quiroga, R. Q. Concept cells: the building blocks of declarative memory functions. Nature Reviews Neuroscience 13, 587–597 (2012).

[11] Mackay, S. et al. Concept and location neurons in the human brain provide the ‘what’and ‘where’in memory formation. Nature communications 15, 7926 (2024).

[12] Kutter, E. F., Bostroem, J., Elger, C. E., Mormann, F. & Nieder, A. Single neurons in the human brain encode numbers. Neuron 100, 753–761 (2018).

[13] Kreiman, G., Koch, C. & Fried, I. Category-specific visual responses of single neurons in the human medial temporal lobe. Nature neuroscience 3, 946–953 (2000).

14. Reber, T. P., et al. Representation of abstract semantic knowledge in populations of human single neurons in the medial temporal lobe. PLoS Biology 17, e3000290 (2019).

[15] Mormann, F. et al. A category-specific response to animals in the right human amygdala. Nature neuroscience 14, 1247–1249 (2011).

[16] Mormann, F. et al. Scene-selective coding by single neurons in the human parahippocampal cortex. Proceedings of the National Academy of Sciences 114, 1153–1158 (2017).

[17] Rey, H. G. et al. Single neuron coding of identity in the human hippocampal formation. Current Biology 30, 1152–1159 (2020).

[18] Reber, T. P. et al. Single-neuron mechanisms of neural adaptation in the human temporal lobe. Nature Communications 14, 2496 (2023).

[19] Cao, R. et al. A neuronal code for object representation and memory in the human amygdala and hippocampus. Nature Communications 16, 1510 (2025).

[20] Hebart, M. N. et al. Things: A database of 1,854 object concepts and more than 26,000 naturalistic object images. PloS one 14, e0223792 (2019).

21. Pilehvar, M. T. & Collier, N. De-conflated semantic representations. *arXiv preprint arXiv:1608.01961* (2016).

[22] Kumar, A. A. Semantic memory: A review of methods, models, and current challenges. Psychonomic bulletin & review 28, 40–80 (2021).

[23] Collins, A. M. & Quillian, M. R. Retrieval time from semantic memory. Journal of verbal learning and verbal behavior 8, 240–247 (1969).

[24] Collins, A. M. & Loftus, E. F. A spreading-activation theory of semantic processing. Psychological review 82, 407 (1975).

[25] Smith, E. E., Shoben, E. J. & Rips, L. J. Structure and process in semantic memory: A featural model for semantic decisions. Psychological review 81, 214 (1974).

26. Hansen, H. & Hebart, M. N. Semantic features of object concepts generated with gpt-3. *arXiv preprint arXiv:2202.03753* (2022).

[27] Tversky, A. Features of similarity. Psychological review 84, 327 (1977).

[28] Wittgenstein, L. Philosophical Investigations (Blackwell, Oxford, UK, 1953). Translated by G.E.M. Anscombe.

[29] Harris, Z. S. Distributional structure. Word 10, 146–162 (1954).

[30] Firth, J. R. Papers in Linguistics, 1934-1951 (Oxford University Press, Oxford, UK, 1957).

[31] Hebart, M. N. et al. Things-data, a multimodal collection of large-scale datasets for investigating object representations in human brain and behavior. Elife 12, e82580 (2023).

[32] Kriegeskorte, N., Mur, M. & Bandettini, P. A. Representational similarity analysis-connecting the branches of systems neuroscience. Frontiers in systems neuroscience 2, 249 (2008).

[33] Miller, G. A. Wordnet: a lexical database for english. Communications of the ACM 38, 39–41 (1995).

[34] Hubel, D. H. & Wiesel, T. N. Receptive fields, binocular interaction and functional architecture in the cat’s visual cortex. The Journal of physiology 160, 106 (1962).

[35] Hubel, D. H. & Wiesel, T. N. Receptive fields and functional architecture of monkey striate cortex. The Journal of physiology 195, 215–243 (1968).

[36] Bitterman, Y., Mukamel, R., Malach, R., Fried, I. & Nelken, I. Ultra-fine frequency tuning revealed in single neurons of human auditory cortex. Nature 451, 197–201 (2008).

37. Kiang, N. Y., Watanabe, T., Thomas, E. C. & Clark, L. F. Discharge patterns of single fibers in the cat’s auditory nerve. (1966).

[38] Kiang, N. & Moxon, E. Tails of tuning curves of auditory-nerve fibers. The Journal of the Acoustical Society of America 55, 620–630 (1974).

[39] Georgopoulos, A. P., Kalaska, J. F., Caminiti, R. & Massey, J. T. On the relations between the direction of two-dimensional arm movements and cell discharge in primate motor cortex. Journal of Neuroscience 2, 1527–1537 (1982).

[40] O’Keefe, J. & Dostrovsky, J. The hippocampus as a spatial map: preliminary evidence from unit activity in the freely-moving rat. Brain research (1971).

[41] Wilson, M. A. & McNaughton, B. L. Dynamics of the hippocampal ensemble code for space. Science 261, 1055–1058 (1993).

[42] Butts, D. A. & Goldman, M. S. Tuning curves, neuronal variability, and sensory coding. PLoS biology 4, e92 (2006).

[43] Shepard, R. N. Toward a universal law of generalization for psychological science. Science 237, 1317–1323 (1987).

[44] Xu, F. & Tenenbaum, J. B. Word learning as bayesian inference. Psychological review 114, 245 (2007).

[45] Quiroga, R. Q. No pattern separation in the human hippocampus. Trends in Cognitive Sciences 24, 994–1007 (2020).

[46] Mikolov, T., Sutskever, I., Chen, K., Corrado, G. S. & Dean, J. Distributed representations of words and phrases and their compositionality. Advances in neural information processing systems 26 (2013).

[47] Boleda, G. & Erk, K. Distributional semantic features as semantic primitives–or not, 2–5 (Stanford, CA, 2015).

48. Park, K., Choe, Y. J., Jiang, Y. & Veitch, V. The geometry of categorical and hierarchical concepts in large language models. *arXiv preprint arXiv:2406.01506* (2024).

[49] Sorscher, B., Ganguli, S. & Sompolinsky, H. Neural representational geometry underlies few-shot concept learning. Proceedings of the National Academy of Sciences 119, e2200800119 (2022).

[50] Ebitz, R. B. & Hayden, B. Y. The population doctrine in cognitive neuroscience. Neuron 109, 3055–3068 (2021).

[51] Franch, M. et al. A vectorial code for semantics in human hippocampus. bioRxiv 2025–02 (2025).

[52] Chang, L. & Tsao, D. Y. The code for facial identity in the primate brain. Cell 169, 1013–1028 (2017).

[53] Leopold, D. A., Bondar, I. V. & Giese, M. A. Norm-based face encoding by single neurons in the monkey inferotemporal cortex. Nature 442, 572–575 (2006).

[54] Quiroga, R. Q., Reddy, L., Kreiman, G., Koch, C. & Fried, I. Invariant visual representation by single neurons in the human brain. Nature 435, 1102–1107 (2005).

[55] Bausch, M. et al. Concept neurons in the human medial temporal lobe flexibly represent abstract relations between concepts. Nature communications 12, 6164 (2021).

[56] Niediek, J., Boström, J., Elger, C. E. & Mormann, F. Reliable analysis of single-unit recordings from the human brain under noisy conditions: tracking neurons over hours. PloS one 11, e0166598 (2016).

[57] Dehnen, G. et al. Duplicate detection of spike events: a relevant problem in human single-unit recordings. Brain Sciences 11, 761 (2021).

[58] Knieling, S. et al. An online adaptive screening procedure for selective neuronal responses. Journal of neuroscience methods 291, 36–42 (2017).

59. Darcher, A., Rapp, R. & Mueller, T. Epiphyte (2024). URL https://github.com/mackelab/epiphyte.

[60] Mormann, F. et al. Latency and selectivity of single neurons indicate hierarchical processing in the human medial temporal lobe. Journal of neuroscience 28, 8865–8872 (2008).

